# Linking coronary microvascular structure and function in preclinical models of coronary microvascular disease

**DOI:** 10.64898/2026.08.31.748256

**Authors:** Mansi B. Kumar, Varun Kanangat, Matthew Woods, Octavio Lopez, Li Li, Eric Blankemeyer, Donna M. Conlon, Uduak George, Scott D. Metzler, Marie A. Guerraty

## Abstract

Despite growing awareness of the importance of the coronary microvasculature in cardiac health and disease, Coronary Microvascular Disease (CMVD) remains poorly understood, underdiagnosed, and without targeted therapies. Preclinical models and quantitative tools to measure CMVD are needed. Here, we use two novel quantitative methods to assess coronary microvascular structure (multi-fractal spectrum analysis) and function (Single Photon Emission Computed Tomography (SPECT)-based intramyocardial blood volume (IMBV) imaging) in mouse models of CMVD. Change in IMBV (ΔIMBV), which serves as a quantitative measure of vasodilatory capacity or microvascular function, was reduced with aging, driven by a significant decrease in female mice. Male mice on ApoE^-/-^ background, fed a high-fat diet (HFD) for 6 months, or both had significantly reduced ΔIMBV. We used immunofluorescence to assess both traditional capillary density and global branching structure and vessel heterogeneity using multifractal spectrum analysis. Both were significantly reduced in all groups, and linear regression modeling showed that they were independently associated with ΔIMBV. Finally, we used ΔIMBV to assess the effects of widely used control Adeno-associated viral (AAV) vectors on coronary microvascular function. AAV-overexpression of GFP did not affect function, but Cre-recombinase compromised coronary microvascular structure and function by 16 weeks. In summary, quantitative assessments of coronary microvascular structure and function highlight changes consistent with CMVD seen with aging, female sex, and metabolic insults. Functional changes are partially driven, but not fully defined, by changes in the underlying structure, highlighting the important and incomplete link between structure and function.

## Introduction

The coronary microvasculature regulates coronary blood flow to meet cardiac metabolic needs. Coronary microvascular disease (CMVD) affects the coronary pre-arterioles, arterioles, and capillaries and disrupts proper matching of blood supply to cardiac demand, leading to ischemia. CMVD may account for as much as 30-50% of the burden of ischemic heart disease, and many individuals with Ischemia or Myocardial Infarction with Non-Obstructed Coronary Arteries (INOCA, and MINOCA, respectively) have CMVD.^1,2^ Risk factors for CMVD include age, hypertension, obesity, hypercholesterolemia, and diabetes, among others.^2,3^ Despite increasing awareness, CMVD remains understudied with limited evidence-based treatments and no directed therapies.^2,4^

To study underlying mechanisms of and evaluate potential therapeutics for CMVD, rigorous, efficient, and quantitative measures of coronary microvascular function and structure in pre-clinical models are needed. In humans, non-invasive imaging methods, such as perfusion PET stress testing,^5,6^ can measure coronary flow and flow reserve (CFR) as measures of microvascular function. Prior groups have used preclinical non-invasive imaging methods to better understand CMVD pathogenesis. For example, Shah et al used cardiac MRI to measure reduced CFR in mice fed a high fat diet,^7^ and Schiattarella et al. used Doppler echocardiography to show reduced CFR in a mouse model of heart failure with preserved ejection fraction due to hypertensive and metabolic insults.^8^ However, these study studies were limited to few cohorts and did not define the link between structure and function.

Previously, we established the feasibility of measuring coronary microvascular function in mice using μSPECT to quantify changes in intramyocardial blood volume (IMBV), as others have in larger animal models.^9,10^ IMBV, defined as the blood contained within the vessels in the left ventricular wall, can be measured using 99mTechnetium (Tc99m)-Pyrophosphate (PYP)-labeled red blood cells (RBC). IMBV increases under hyperemic conditions, reflecting coronary vasodilatory capacity and microvascular function. Here, we optimized our μSPECT methods for higher throughput imaging and assessed the effects of known CMVD risk factors, including aging, sex, metabolic dysfunction, and hypercholesterolemia, on coronary microvascular function in mouse models. We compared microvascular function with quantitative measures of coronary microvascular structure using both traditional capillary density and more holistic multifractal assessment. Finally, we used these tools to evaluate the effects of standard Adeno-Associated Virus (AAV)-based conditions on coronary microvascular structure and function.

## Methods

### Mouse models

Our study examined male and female animals, and sex-dimorphic effects are reported. An overview of the study methodologies and mouse cohorts is presented in Figure 1. Eight week old C57BL/6J mice and Apolipoprotein E knockout (ApoE^-/-^) mice were purchased from Jackson Laboratories (Bar Harbor, ME). All mice were housed 3-5 mice/cage in a climate-controlled room with a 12:12 light:dark cycle with *ad libitum* access to regular chow diet. Where specifically noted, animals were fed *ad libitum* a high fat diet (HFD) containing 45% kcal of fat (D12451, Research Diets) for a duration of 6 months. Imaging studies were performed on 10-35 week old male and female mice. Mice were euthanized by cervical dislocation after 4% isoflurane inhalation. All studies were approved by the University of Pennsylvania Institutional Animal Care and Use Committee and conform to the NIH Guide for the Care and Use of Laboratory Animals.

**Figure 1.**
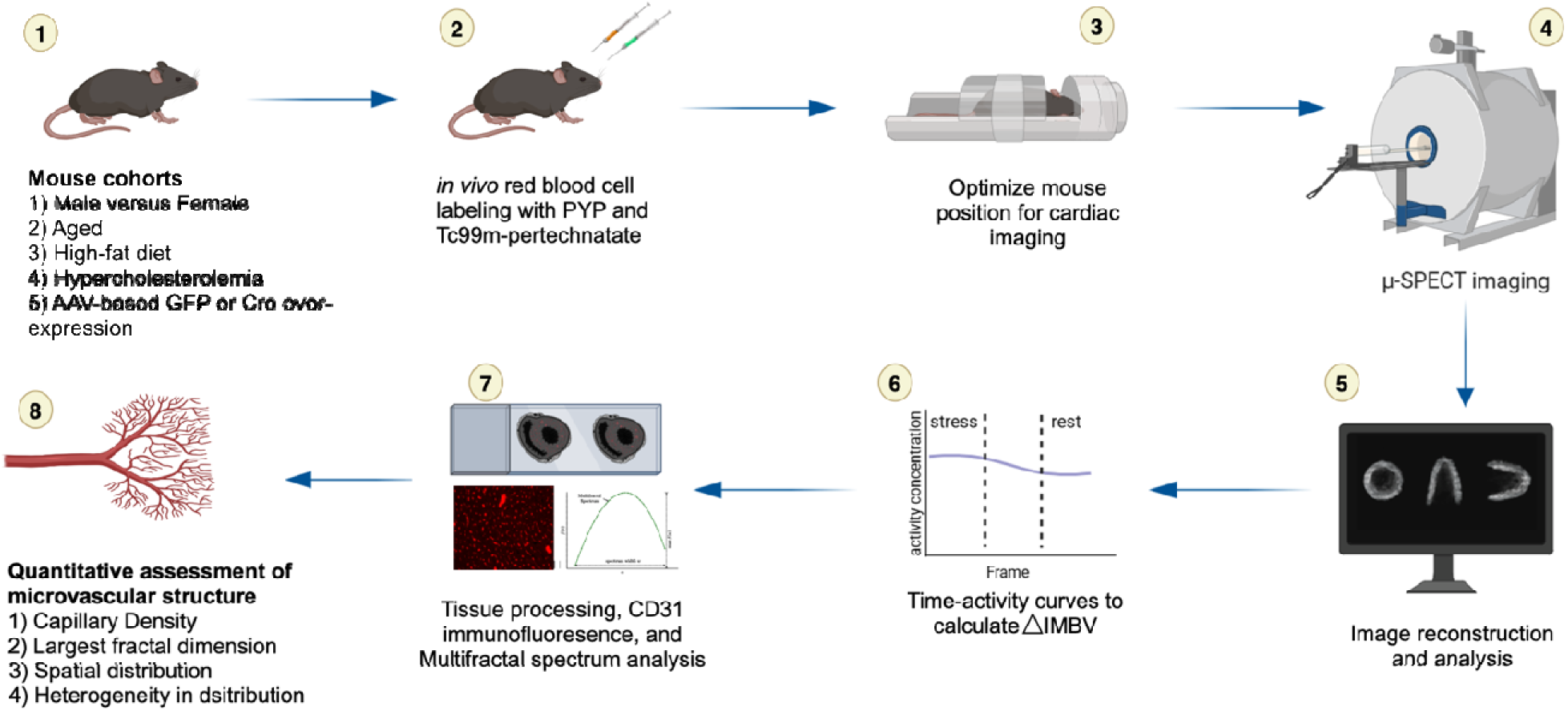
Overview of workflow including experimental conditions, image acquisition, and tissue processing. (1) Distinct cohorts of mice treated with CMVD risk factors or common control AAV vectors underwent RBC labeling (2) followed by SPECT imaging (3,4). Image reconstruction and analysis (5) generates time activity curves acquired under hyperemic (stress) and basal (rest) to calculate IMBV. (7) Tissue is then processed for histology, and vessels are visualized using CD31 immunofluorescence. (8) Capillary density and several multifractal parameters are obtained. Figure made with Biorender.

### AAV injection

Adeno-associated virus (AAV)-based cardiac-specific overexpression is a widely used tool in cardiovascular research. We therefore assessed the effects of two commonly used AAV-based control vectors (AAV9.cTnT.PI.eGFP and AAV9.cTnT.PI.iCre.IRES.tdTomato) on coronary microvascular structure and function. 8-10 week old C57BL/6J mice were injected with 3x10^12 viral particles of AAV9-GFP or 5x10^11 viral particles of AAV9-Cre.^11^ All AAVs were synthesized by the University of Pennsylvania Vector Core.

### IMBV image acquisition

*In vivo* RBC labeling with Tc99m-PYP has been previously described.^9^ Briefly, mice were injected with 100 μL commercial PYP (Pharmalucence, Billerica, MA) reconstituted in saline, and, following a 20-minute delay, injected with 4-6 mCi of ^99m^Tc-Pertechnatate. Mice were sedated with 2.5% isoflurane and imaged with the U-SPECT instrument (MI labs, The Netherlands) for 5 minutes using one-minute frames in 3 bed positions. Isoflurane was then reduced to 1.25% and then mice were imaged for an additional 8-10 minutes. In a subset of mice, we evaluated the effects of 3% compared to 2.5% isoflurane as a hyperemic condition. Basal conditions were assessed under 1.25% isoflurane.^12^

### Image analysis

Acquired images were reconstructed using 0.2mm cubic voxels with 10 iterations and 4 subsets and then post-filtered (0.25 mm FWHM Gaussian filter). We used an in-house developed MATLAB software to obtain cardiac short axis, horizontal long axis (HLA), and vertical long axis (VLA). We customized regions of interest (ROIs) to fit the myocardium and blood pool of each mouse and then extracted the time activity concentration curves. For each time-activity curve, we averaged the first four IMBV values as our IMBV_stress_ and the last four IMBV values as our IMBV_rest_ (Supplemental Figure 1).

### Mixing parameters and ***Δ***IMBV

To account for mixing between blood pool and myocardial ROIs, we previously defined mixing parameters which represent the spillover activity from blood pool to LV and LV into blood pool (F_A_ and F_T_, respectively), and recovery coefficients of the myocardium and blood pool (R_T_ and R_A_, respectively).^13^ Here, we set R_A_ and R_T_ as 1, F_T_ as 0.29, and F_A_ as 0.25. Using these parameters as coefficients we calculated the relative change in IMBV based on the ratio of IMBV_stress_ to IMBV_rest_ (Equation 1), where *θ* = ratio of measured blood pool activity under rest and stress; *η* = ratio of measured LV myocardium activity under rest and stress:

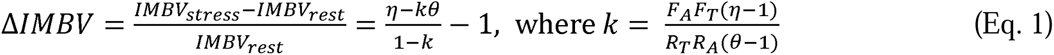

### Tissue processing and capillary density

We next evaluated capillary density in the hearts of the imaged mice. Briefly, hearts were collected from a subset of imaged mice and processed for histology. Tissue was fixed in 4% formaldehyde, embedded in paraffin, and sectioned into 6um sections. To visualize capillaries the slides were stained with rat anti-mouse CD31 (1:200, Biozol Diagnostics) and ImpressRed secondary antibody (1:1000, Vector Labs) and imaged on a Keyence microscope (BZ-X1000). Nine high power images (40x) were acquired per mouse, three each from the epicardial, mid myocardial, and endocardial regions of left ventricle (LV) by an operator blinded to condition. Image J was used to quantify capillary density. We normalized the non-quantified regions by dividing the counted capillaries by the percentage of quantified area [11, 12]. Data are presented as number of capillaries per high-powered field (#/hpf).

### Multifractal analysis

To more fully capture coronary structure and rigorously assess for multiscale differences, we used multifractal spectrum analysis. Multifractal theory is a mathematical framework that is often used to describe and analyze complex branching structures that vary across multiple spatial scales.^14,15^ A key concept in multifractal theory is the multifractal spectrum *f*(*α*), a concave down function, that captures the scaling (i.e., the spatial differences in the distribution of structural features).^16^ Briefly, the coronary microvasculature was segmented from the immunofluorescence images, converted to grayscale, binarized, and then subsequently used to compute the multifractal spectrum. Here we consider three key variables of *f*(*α*) (Figure 1): (1) maximum of the *f*(*α*) function, which represent the largest fractal dimension, (2) the *α* at which that maximum is found (i.e. *a*rgmax for *f*(*α*), represents a measure of how spatially distributed the coronary microvasculature are, and (3) the width of the multifractal spectrum, which reflects how unevenly the coronary microvasculature are distributed, whether they are arranged in a more uniform pattern or vary greatly from one tissue region to another. Please see Supplemental Methods for additional details on the multifractal spectrum computation.

### Statistical analysis

Data were plotted as mean with standard deviations or median with inter-quartile range as indicated. Outliers were excluded using Rout’s outlier test with Q=2%. Differences in the mean values between 2 groups were assessed using 2-tailed Student’s *t* test. Differences in mean values among more than 2 groups were assessed by 1-way ANOVA with any individual group differences determined using Tukey’s test and p-value < 0.05 was considered significant. Least squares multiple linear regression analysis was used for multivariate analysis to compare measure of structure and function. To compare male and female ΔIMBV, we used 2-way ANOVA with Sidak’s test for multiple comparisons. All tests were performed using Prism 10.4.0.

## Results

### IMBV Imaging

We have previously reported the use of μSPECT to quantify intramyocardial blood volume.^9^ To allow for higher throughput imaging and minimize inter-animal variability, we optimized our previously reported IMBV protocol (Figure 1). Briefly, we performed *in vivo* RBC labelling using Tc99m, acquired 1-minute frames and used an in-house developed MATLAB software to visualize the three orthogonal cardiac planes and left ventricular myocardial and blood pool regions of interest (ROIs) (Supplemental Figure S1A,B). Average LV and blood pool uptake were obtained under hyperemic (stress) and basal (rest) conditions and used to calculate IMBV as outlined above (Equation 1). We next assessed the effects of 2.5% versus 3% isoflurane as hyperemic conditions and found no difference (Supplemental Figure S2A, B). We therefore used 2.5% isoflurane to induce hyperemia for all subsequent imaging studies and used 1.25% isoflurane as the basal or rest condition (Supplemental Figure S1C). These methods showed minimal inter-observer variability (Supplemental Figure S2C). Finally, to compensate for motion during imaging protocol, we found that frame-by-frame ROI analysis reduced variability compared to using a summed image of all the frames (Supplementary Figure 2D,E). With these modifications, we were able to reliably quantify ΔIMBV in larger cohorts of mice.

### The effects of CMVD risk factors on ***Δ***IMBV

We imaged male and female wildtype mice aged 8 weeks to establish baseline ΔIMBV of 29 ± 5.7% (n = 7 each male and female mice). When aged to 9 months, mice had reduced ΔIMBV of 20 ± 8.9% (p<0.01, n=9 each males and females, Figure 2A). Interestingly, this signal was primarily driven by a reduced ΔIMBV in female mice (29.9 ±6.3% vs 17.1 ± 7.9%, p=0.005), though male mice showed a non-significant trend (22.8 ± 5.2% vs 22.8 ± 6.3% p=0.18). We did not observe sex-based statistically significant differences across control mice or the aged mice (Figure 2B).

**Figure 2.**
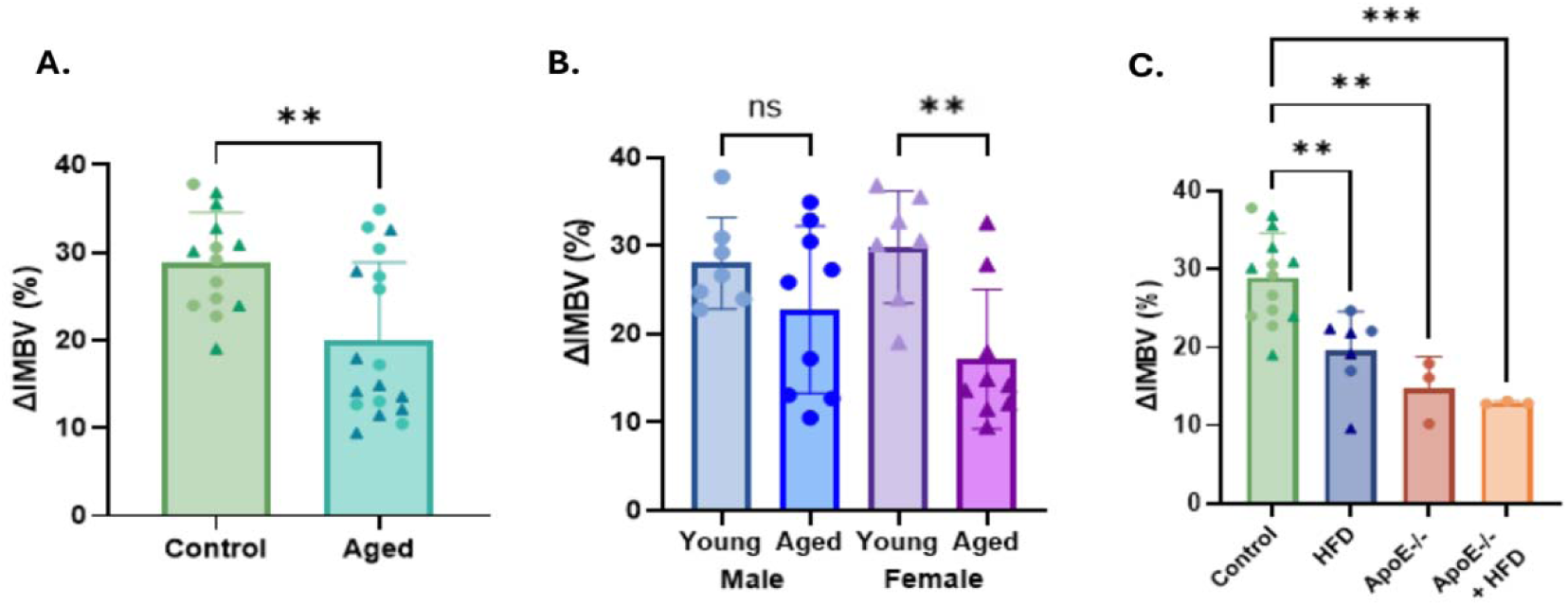
ΔIMBV across different cardiovascular risk factors. (A). ΔIMBV was reduced in mice aged for 10 months on chow diet relative to 8 week control mice by Student’s t-test. (B) There is no sex difference in ΔIMBV at baseline (n = 7 each male and female mice) Aging reduced ΔIMBV in female mice, but the effect was blunted in male mice. (n=9 each males and females) by 2-way ANOVA (C) 6 months of high fat diet (HFD) or loss of ApoE to induce hypercholesteremia significantly reduces ΔIMBV (n=14, 7, 3 and 3 respectively). Triangles represent female mice and circles represent male mice. All groups compared to control by ANOVA with multiple comparisons. **p<0.01,

In addition to age, metabolic insults are known risk factors for CMVD. Indeed, mice maintained on a 45% high fat diet (HFD) for 6 months had a significant 0.69-fold reduction in ΔIMBV (19.5±5.0%, p<0.01). Similarly, ApoE^-/-^ mice, which develop hypercholesterolemia over time,^17^ showed a similar reduction in ΔIMBV whether maintained on chow diet (14.8±4.0%, p<0.01) or HFD (12.9±0.2%, p<0.001) (Figure 2C). Both groups had similarly elevated total cholesterol levels (452 ± 312 mg/dl vs 444 ± 167 mg/dl, p = 0.99). Of note and consistent with CMVD, aged, HFD, and ApoE-/- mice fed a HFD all had normal left ventricular ejection fraction (Supplemental Figure S3).

### Capillary density is reduced with metabolic insults and correlates moderately with ***Δ***IMBV

Coronary microvascular disease is a heterogenous process, which includes structural and functional changes to the coronary microvasculature.^18^ Capillary rarefaction represents one structural change often seen in CMVD which can affect coronary flow. We assessed capillary density in our models by quantifying the number of CD-31 positive vessels in fixed hearts after imaging (Figure 3A). Aged mice had no decrease in capillary density relative to controls cohort. However, those maintained on a HFD had significantly reduced capillary density 0.87-fold. ApoE-/- mice on either chow or HFD also had decreased capillary density 0.68-fold and 0.57-fold (Figure 3B). We next examined the relationship between our functional microvascular assessment, ΔIMBV, with capillary density. Across all mice, there is a significant linear relationship with moderate R^2^ suggesting capillary density only partially accounts for vasodilatory capacity (R^2^=0.25, p<0.01) (Figure 3C).

**Figure 3.**
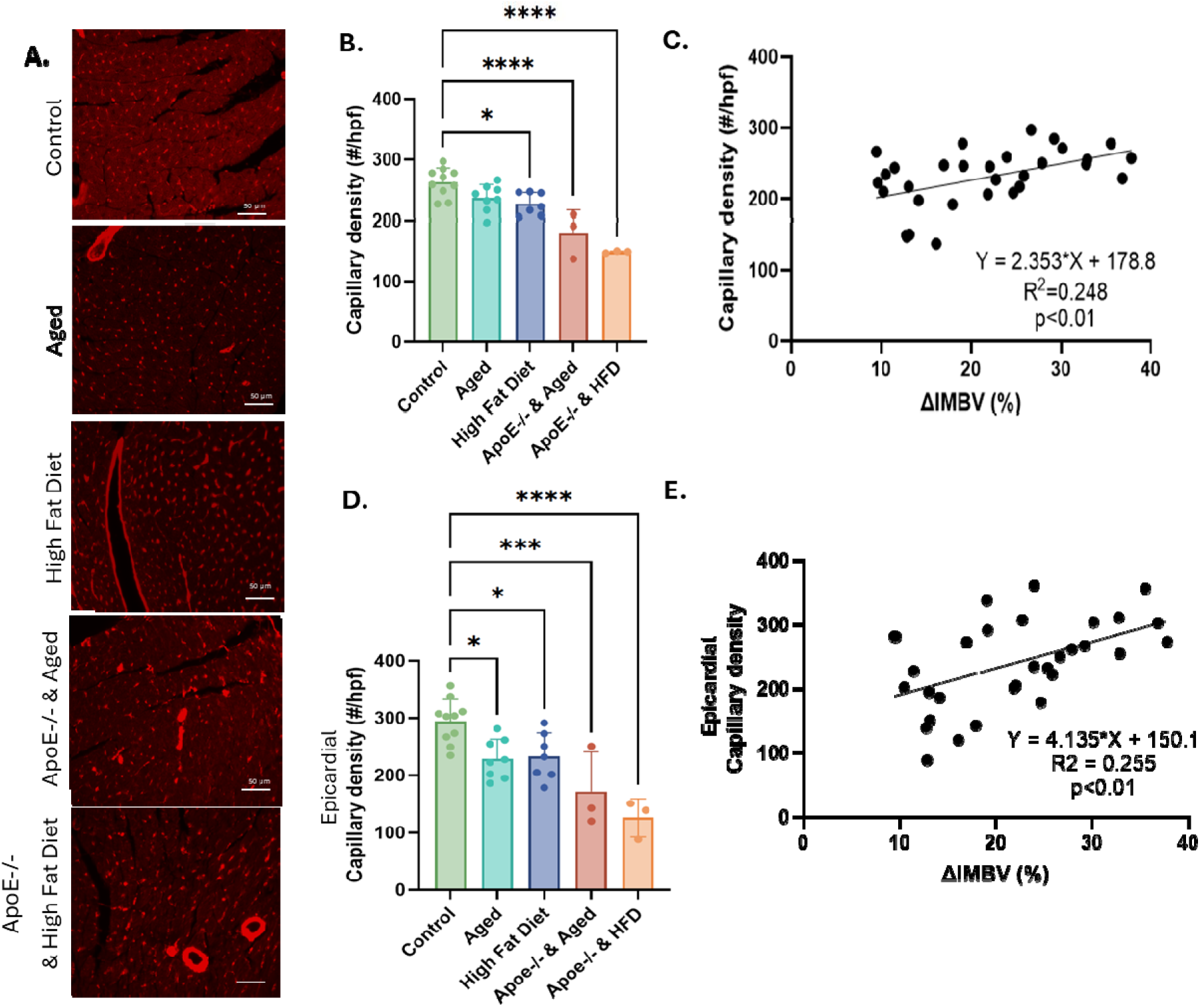
Capillary density is reduced with metabolic insults and correlates with ΔIMBV. (A) Representative images high power fields showing CD31 staining. (B) Capillary density was decreased in high fat diet and ApoE-/- mice. Data represents mean quantification of 9 images per mouse, n=3-16 per group. All groups compared to control by ANOVA with multiple comparisons Data presented as number capillaries per high-[owered field (#/hpf) (C) Across all mice, there is a significant linear relationship between ΔIMBV and capillary density. (D,E) Changes in capillary density across groups and linear relationship between capillary density and IMBV were mirrored in the epicardial samples. Data represents mean quantification of 3 images per mouse, n=3-16 per group *p<0.05, ***p<0.001, ****p<0.0001.

Capillary density is measured by sampling diffusely throughout the myocardium. However, differences in microvascular distribution have been reported.^19^ We therefore assessed capillary density in the epicardium (epi), mid-myocardium (mid), and endocardium (endo). We found that changes in capillary density were driven largely by changes in the epicardium (Figure 3D, Supplemental Figure S4). In particular, capillary density in the epi region decreased with aging by 0.77-fold (229.2 ± 34.4 #/hpf versus 294.8±38.1#/hpf, p<0.5, Figure 3D), serving as a more sensitive measure than global capillary density. Mid and endo capillary density was only reduced in ApoE-/- models (Supplemental Figure S4A,B), and there was no relationship between mid and endo capillary density and ΔIMBV (Supplemental Figure S4C,D).

### Multifractal analysis provides independent and additive structural information

While capillary density is often used to assess the coronary microvasculature, CMVD is associated with other vascular changes, such as increased arteriolar size heterogeneity or loss of homogeneous vessel distribution. These are more difficult to assess, though new quantitative tools have emerged. For example, multifractal spectrum provides a more holistic assessment of changes in vessel size, branching, and spatial distribution. It uses a concave down curve to summarize several parameters that reflect properties of a branching structure. In particular, α is plotted on the x-axis and *f*(*α*), which represents the corresponding fractal dimension, is plotted on the y-axis. The most prevalent fractal seen in the structure is the peak of the curve (i.e. argmax *f*(*α*), termed argmax herein). Spectrum width reflects the heterogeneity of branching distribution. Finally, symmetry of the curve also informs on which aspects of branching may be missing. A representative multifractal spectrum from a CD31-stained myocardial section is shown in Figure 4A. We found that multifractal parameters, while distinct from capillary density, were also associated with IMBV. There was a statistically significant linear relationship between argmax and ΔIMBV (R^2^=0.24, p<0.01) (Figure 4B).

**Figure 4.**
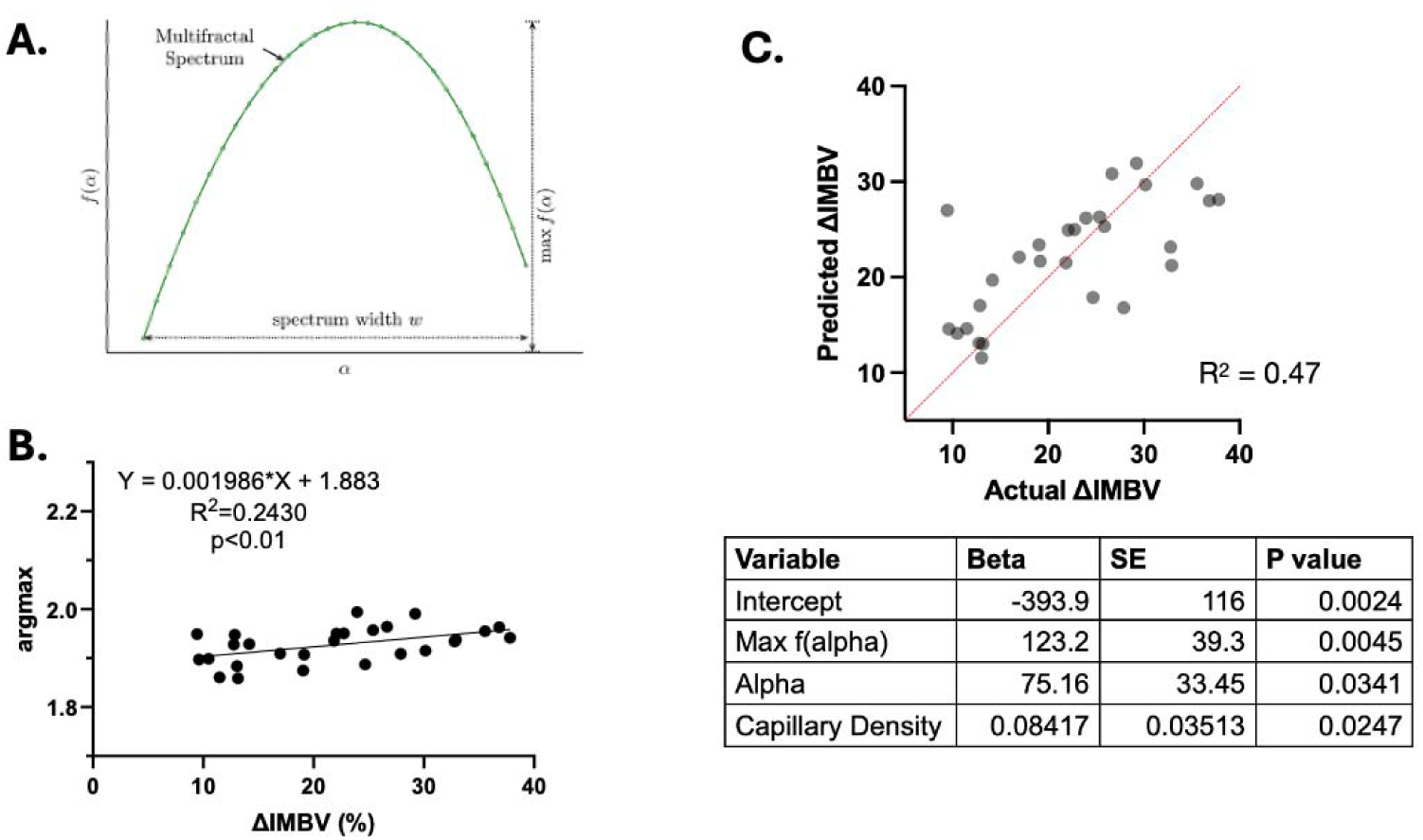
Multifractal spectral analysis captures independent structural parameters associated with ΔIMBV. (A) Schematic of a multifractal spectrum showing the max and the spectrum width noted. Arg max for is the value of at which attains its maximum. (B) Scatterplot highlighting significant linear relationship between argmax and ΔIMBV. (C) Linear regression modeling showing that arg max (Alpha), max, and capillary density are independently associated with ΔIMBV.

To determine whether multifractal spectrum provides additional structural data over traditional capillary density, we performed regression modeling using capillary density and multifractal parameters. We found that max *f*(*α*) and argmax *α* were independently associated with IMBV and provided additional structural data beyond capillary density. In particular, combining multifractal with capillary density data increased the R^2^ from 0.24 to 0.47 (Figure 4C), suggesting that almost half of the variability noted in IMBV can be modeled with these structural parameters.

### Effect of AAV9 on IMBV and capillary density

Viral-based vectors have emerged as a useful tool to assess the effects of genes *in vivo,* and AAV-mediated overexpression is often used to interrogate changes over time.^20,21^ We therefore assessed the effects of two commonly used AAV-controls (AAV9-GFP and AAV9-Cre) on coronary microvascular structure and function. We found AAV9-GFP did not affect ΔIMBV, as compared to the control group in Figure 2 (Figure 5A), and capillary density was also unchanged 4 weeks post-injection in female mice injected with 3e12 vp (Figure 5B). Wildtype littermate control mice were injected with 5e11 vp AAV9-Cre and μSPECT imaging was performed on these mice 5 weeks post-injection. There was no significant difference in ΔIMBV as compared to the AAV9-GFP mice (Figure 5A). To determine if longer cardiomyocyte exposure to Cre expression affected IMBV, we injected a separate cohort of mice with 5e11vp of AAV9 Cre and imaged the mice 8 and 16 weeks post-injection. While there was still no difference in ΔIMBV after 8 weeks, there was a 50% decrease in ΔIMBV 16 weeks post-injection (30.3 ± 1.4% vs 15.4 ± 3.4%, p<0.01 n=5 females per group Figure 5A). In addition, capillary density was decreased 37% with mice injected with AAV9-Cre after 16 weeks as compared to 5 weeks (308.9 ± 27.8 #/hpf, 196.4 ± 22.1 #/hpf, p<0.001, Figure 5B-D). Overall, capillary density and ΔIMBV have a significant linear relationship with AAV groups (R^2^=0.31, p=0.0003) (Figure 5E).

**Figure 5.**
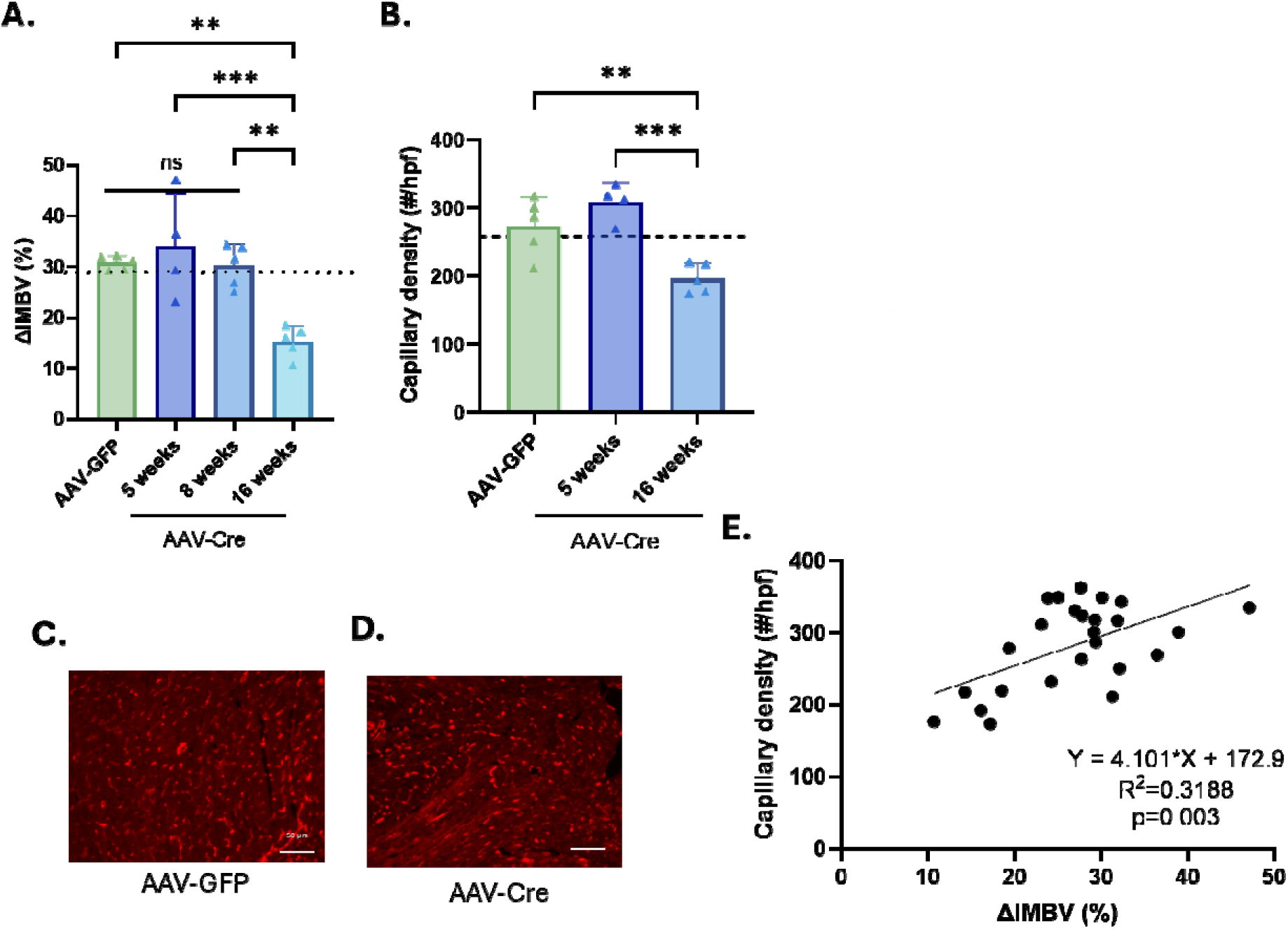
Assessment of AAV on ΔIMBV and capillary density. (A) AAV9-GFP and AAV9-Cre after 5 or 8 weeks do not impact ΔIMBV while AAV9-Cre decreases ΔIMBV 16 weeks post injection. n=10, 13, and 7 respectively (B) AAV9-GFP does not impact capillary density but AAV9-Cre reduces capillary density significantly after 16 weeks. Data represents the mean quantification of 9 images per mouse, n= 10,13, and 7, respectively. All groups compared by ANOVA with multiple comparisons. Dotted line represents control wildtype mouse values presented in Figure 2A and 3B respectively Data presented as number capillaries per high-powered field (#/hpf) (C,D) Representative images of capillary density for AAV9-GFP, and AAV9-Cre groups. (E) There is a significant linear relationship between capillary density and ΔIMBV after AAV treatment. **p<0.01, ***p<0.001, scale bar = 50um.

There were also regional differences in capillary density noted. In contrast to the aged and metabolic dysfunction models, AAV9-Cre reduces capillary density uniformly throughout the epi, mid, and endo regions, as compared to both AAV9-GFP and AAV9-Cre 5 weeks post injection (Supplemental Figure S5A-C). However, epi capillary density, but not mid and endo, was significantly associated with ΔIMBV (R^2^=0.28, p=0.006, Supplemental Figure 5D-F).

## Discussion

Here, we report the relationship between μSPECT-based IMBV imaging, as a measure of coronary microvascular function, and both capillary density and multi-fractal spectrum analysis, as measures of structure, in preclinical mouse models of CMVD. We used these quantitative tools to show reduced ΔIMBV with aging in a sex-dependent manner and with metabolic insults. We then compared coronary microvascular function with both traditional capillary density and novel multifractal analysis to show that structure influences, but does not fully define, function. Finally, we used IMBV imaging to show that widely used AAV-control vectors have differential effects on coronary microvascular function.

CMVD is a heterogeneous pathology characterized by both functional and structural changes. These are interrelated as structural changes, when sufficiently severe, can impact vessel function and resting flow. Coronary microvascular function refers to the ability of the microvasculature to appropriately vasodilate and match coronary blood flow to meet cardiac metabolic demand, and dysfunction suggests an impaired ability to augment flow. This may be due to increased resting coronary flow or to reduced hyperemic flow, and underlying structural abnormalities can affect either. For example, under basal conditions, arterioles are the highest resistance part of the coronary microvasculature.^22^ During hyperemic conditions, arteriolar resistance decreases, and increased intravascular pressure is propagated down to the capillaries.^22^ Reduced capillary density may therefore have minimal effect of resting blood flow but could blunt hyperemic flow. Using multifractal analysis, we identified other structural measures which, independent of capillary density, predict ΔIMBV. This further highlights the complex, and incomplete, interplay between coronary microvascular structure and function.

Our findings are consistent with prior preclinical studies showing that cardiovascular risk factors can affect coronary microvascular structure and function.^23,24^ Wang et al. found that a hypercholesteremic Paigen diet for 8 weeks reduced coronary blood flow in mice.^25^ Similarly, others have shown that mice aged 12-18 months or fed HFD for 14 weeks have decreased capillary density compared to young mice.^26,27^ Porcine models of diet-induced hypercholesterolemia and CMVD, respectively, also showed regional changes in microvascular structure differences and reduced coronary flow reserve.^24,28^ Our study also found that epicardial capillary density was a more sensitive measure of microvascular structural changes in mice. In humans and pigs, the sub-endocardium has higher capillary density,^29^ whereas in rodents, capillary density was higher in the epicardial region of the myocardium.^19^

Coronary microvascular dysfunction is reportedly more prevalent in women, however, some imaging studies show similar prevalence between men and women.^3,30,31^ We found no differences in ΔIMBV between young male and female mice. However, the reduction in ΔIMBV seen with aging was sex-dependent and driven by more significant decreases in ΔIMBV in female mice, suggesting a novel interaction between aging and sex in CMVD. Genetic studies have also identified sex-heterogeneity in microvascular function.^32^ Our data are consistent with these findings and suggest that there may be a complex interplay between sex and cardiovascular risk factors. Future studies will be crucial to tease these apart in the context of CMVD.

AAV vectors have emerged as useful tools used to evaluate gene function *in vivo*. We evaluated the effects of common AAV vectors on IMBV and found that longer-term treatment with AAV9-Cre (16 weeks) reduced both capillary density and ΔIMBV. These suggest an ideal timeframe of 5-15 weeks post injection for cardiac-specific Cre-based studies. AAV9-GFP did not affect microvascular structure or function at 4 weeks, and additional studies are needed to assess the long term effects.

In summary, ΔIMBV and multifractal spectrum analysis are technically feasible quantitative approaches to measure coronary microvascular function and structure in mice, and they could be used to assess mechanisms of CMVD and evaluate potential therapies. There are several important limitations. First, we consider coronary microvascular function homogeneously throughout the left ventricle. This is a reasonable assumption for CMVD, but other methods are needed to assess regional flow for non-homogeneous processes, e.g. hypertrophic cardiomyopathy. Second, we use isoflurane 1.25% and 2.5% for rest and hyperemic flow, respectively, as these doses have been shown have minimal effect on heart rate while increasing myocardial blood flow.^12^ Other pharmacologic agents such as dobutamine or dipyridamole can be used, but these require additional sedation and parenteral access. Third, while we obtained serial measures in AAV9-Cre mice showing a change in ΔIMBV, most cohorts did not have sequential imaging to allow for structure-function correlation. Additional time-course studies will be needed. Fourth, we use histology to quantify structure and show that multi-fractal spectral analysis is able to measure branching and higher level structure from a 2D image. Validating this with 3D imaging, including micro-CT (which can provide *in vivo* measurements but fails at the spatial resolution of capillaries) and light-sheet microscopy will establish the ability to measure 3D branching structure. Similarly, comparing IMBV imaging with microvascular function assessments by ultrasound and cardiac MR will also be an important future direction. Further studies are also needed to link coronary microvascular structure and function in larger animal models and in humans to fully elucidate these relationships. Finally, while IMBV can be used as a high throughput and robust approach to measure coronary microvascular function, and may inform on resting and hyperemic conditions separately, additional methods to measure absolute myocardial blood flow quantification will be helpful to further assess mechanism of CMVD.

## Author Contributions

MK, VK, EB, OL, LL, MW, DMC, UG, SM performed experiments and data analysis. MK, VK, MAG, and UG wrote the manuscript. MW, OL, EB, DMC, UG, SCM, MAG edited the manuscript. UG, MAG, and SDM obtained funding and supervised the work.

## Acknowledgements

We acknowledge the University of Pennsylvania Small Animal Imaging Facility for their support with imaging and instrumentation.

