## Supplemental Figures for "Linking coronary microvascular structure and function in preclinical models of coronary microvascular disease"

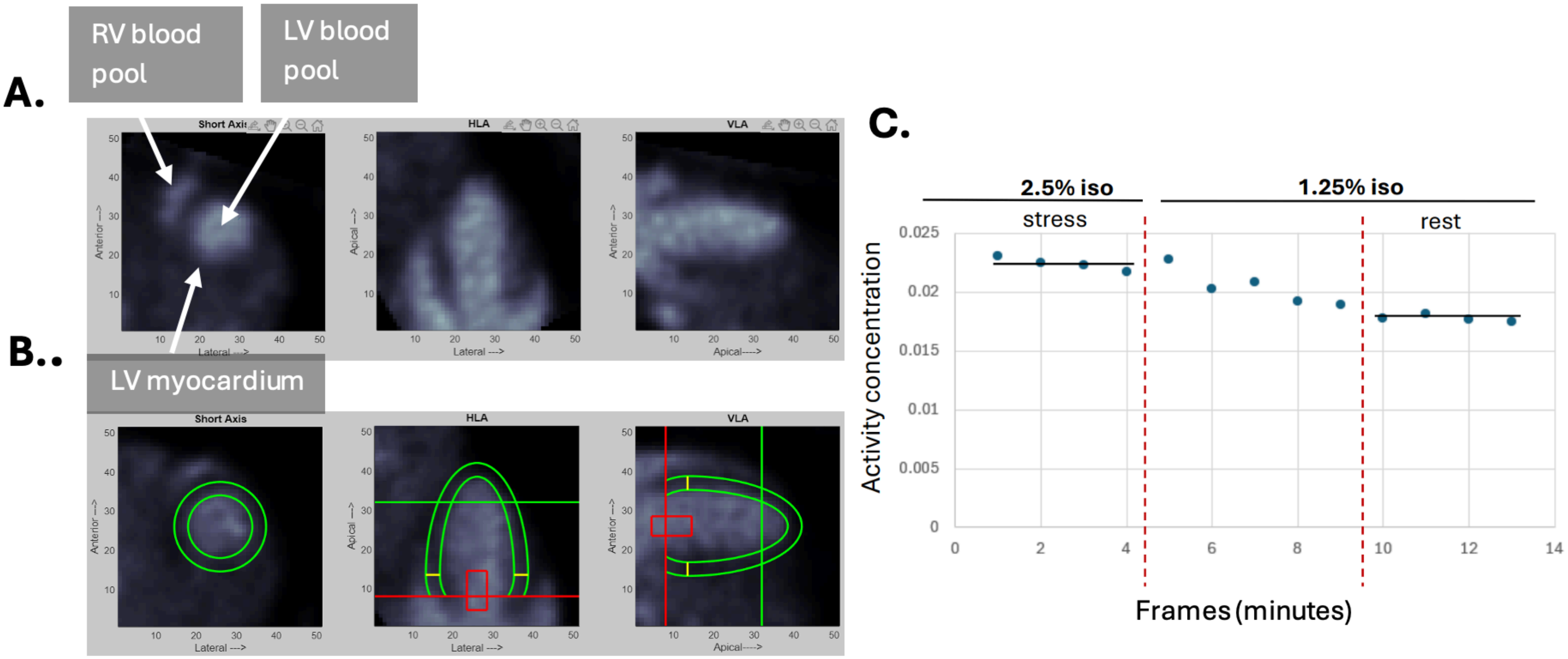


**Supplemental Figure S1. Representative images and time-activity curves.** (A) Orthogonal short axis, horizontal long axis (HLA), and vertical long axis (VLA) images of mouse heart with labelled RBCs (white) and dark tissue. (B) LV myocardial (green) and blood pool (red box) regions ROIs for which activity concentration is measured. Red line = mitral valve plane. (C) Representative time activity curve for LV myocardial ROI. Activity is higher under hyperemic conditions (2.5% isoflurane) and then decreases under basal conditions (1.25% isoflurane). A Washout period is demarcated by vertical dashed lines.

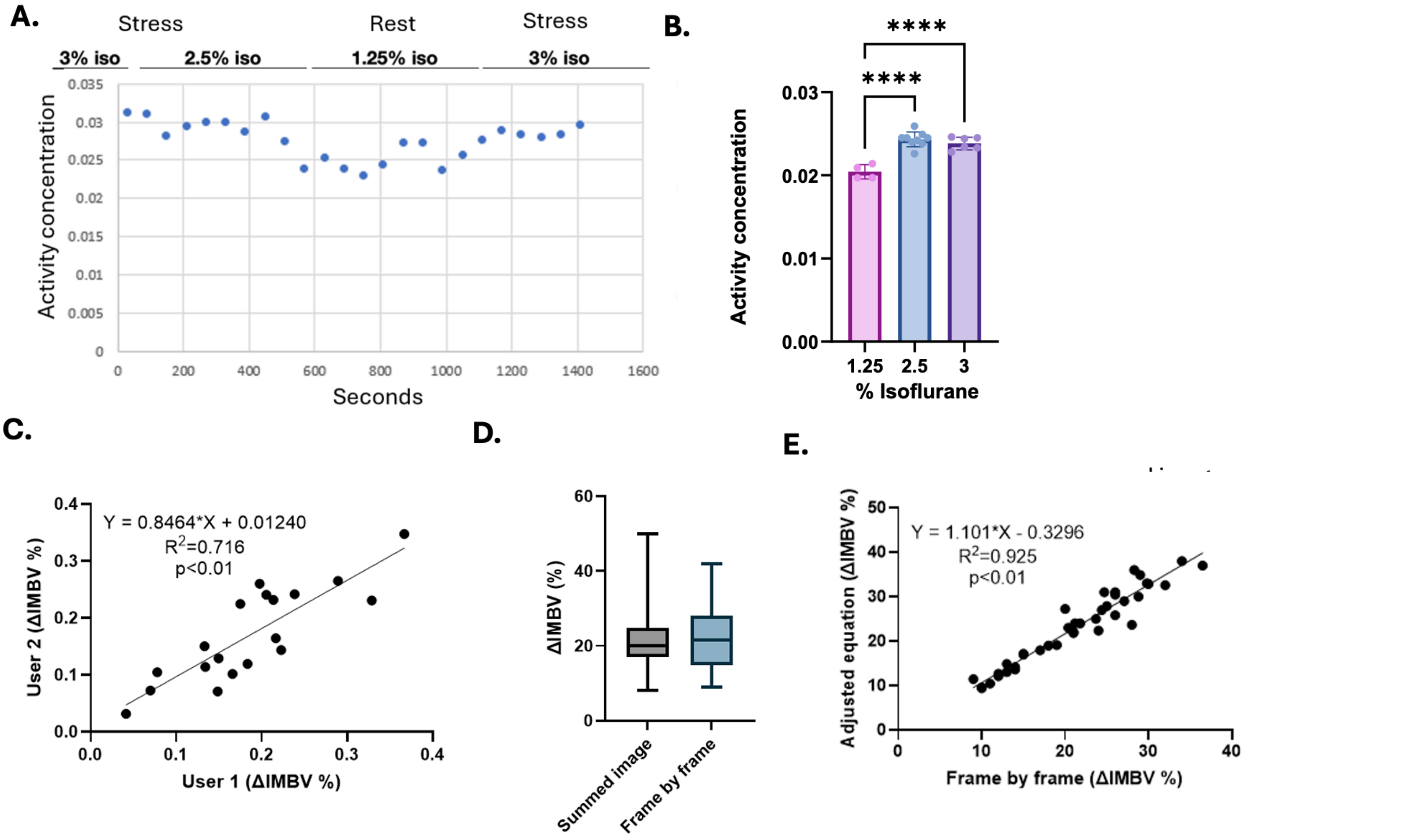


**Supplemental Figure S2. IMBV optimization.** (A) Representative time-activity curve comparing 2.5% isoflurane and 3% isoflurane for hyperemic stress condition. (B) Activity concentration in the myocardial ROI is lower during rest and increases with 2.5% and 3% isoflurane, with no difference between 2.5% and 3% isoflurane. n=4, 9, and 6 mice per group, respectively. All groups compared to control by ANOVA with multiple comparisons, ****$p<0.0001$. (C) Scatterplot showing user inter-user variability in quantifying IMBV, n=19. (D) Drawing ROIs frame-by-frame rather than a single time for the summed image reduced variability without changing the mean IMBV. (E) Adjusted equation 1 for calculating IMBV is similar to measuring IMBV as the % difference between rest and stress, n=33.

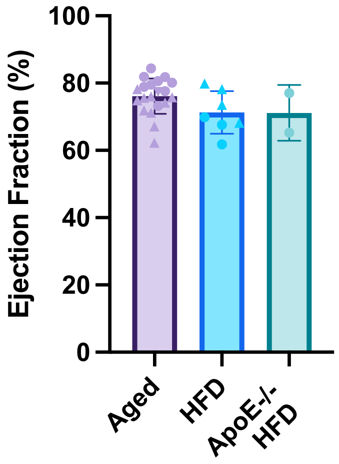


**Supplemental Figure S3. Ejection Fraction assessed by echocardiogram.** Ejection fraction assessed by 2D echocardiography was normal for aged, HFD, and ApoE-/- mice on HFD. n = 20, 7, and 2 mice per group, respectively.

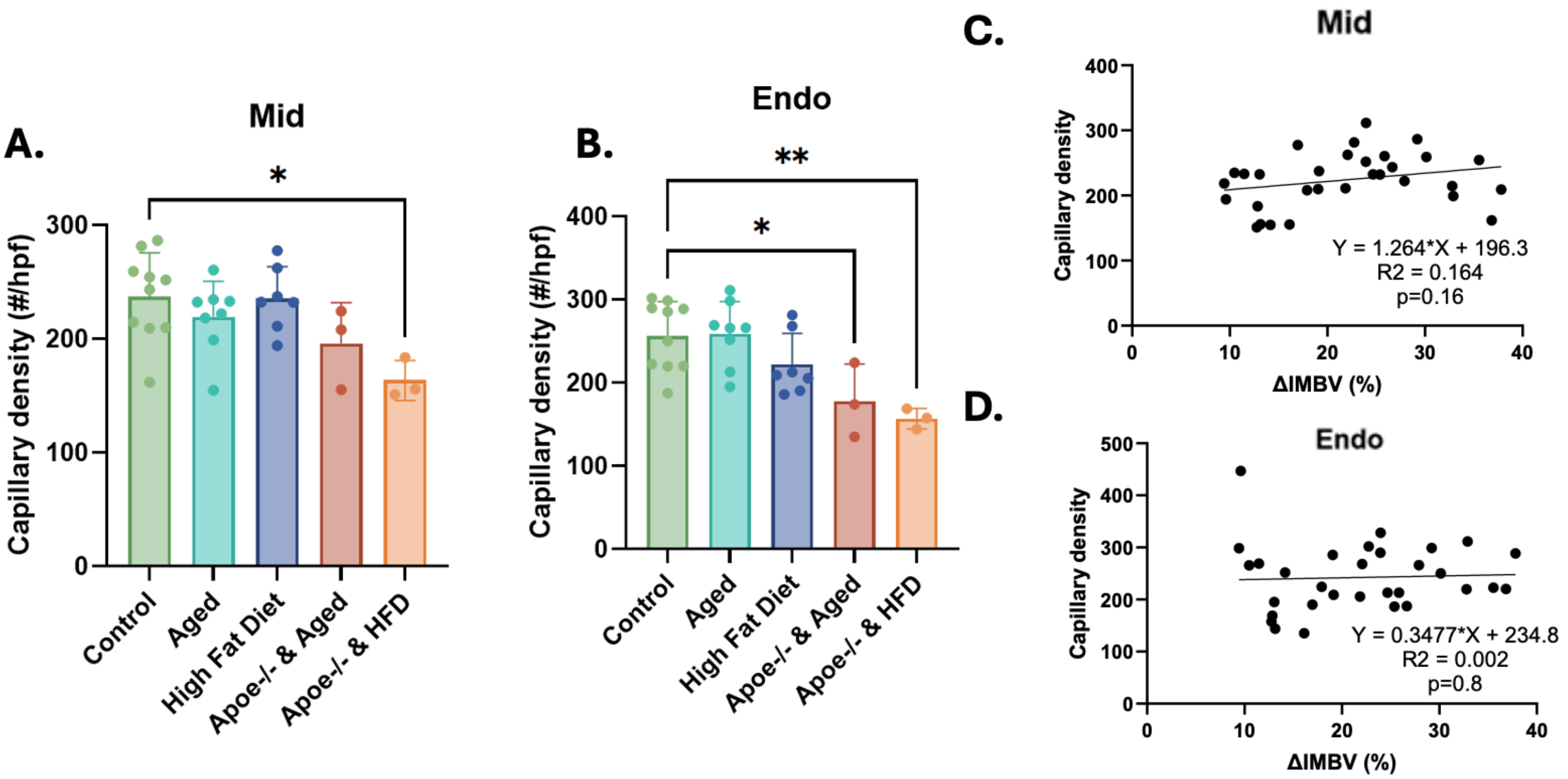


**Supplemental Figure S4. Capillary density across the mid-myocardial and endocardial regions.** (A) Capillary density in the mid-myocardial region was was the least density and decreased only in ApoE-/- & HFD groups. (B) Endocardial capillary density was reduced in both ApoE-/- groups. (C, D) There was no significant relationship between regional capillary density and IMBV for the mid-myocardial and endocardial regions. Data represent mean quantification of 3 images for each of 3 regions per mouse, n=10, 8, 7, 3, and 3 per group, respectively. All groups compared by ANOVA with multiple comparisons, *p<0.05, **p<0.01.

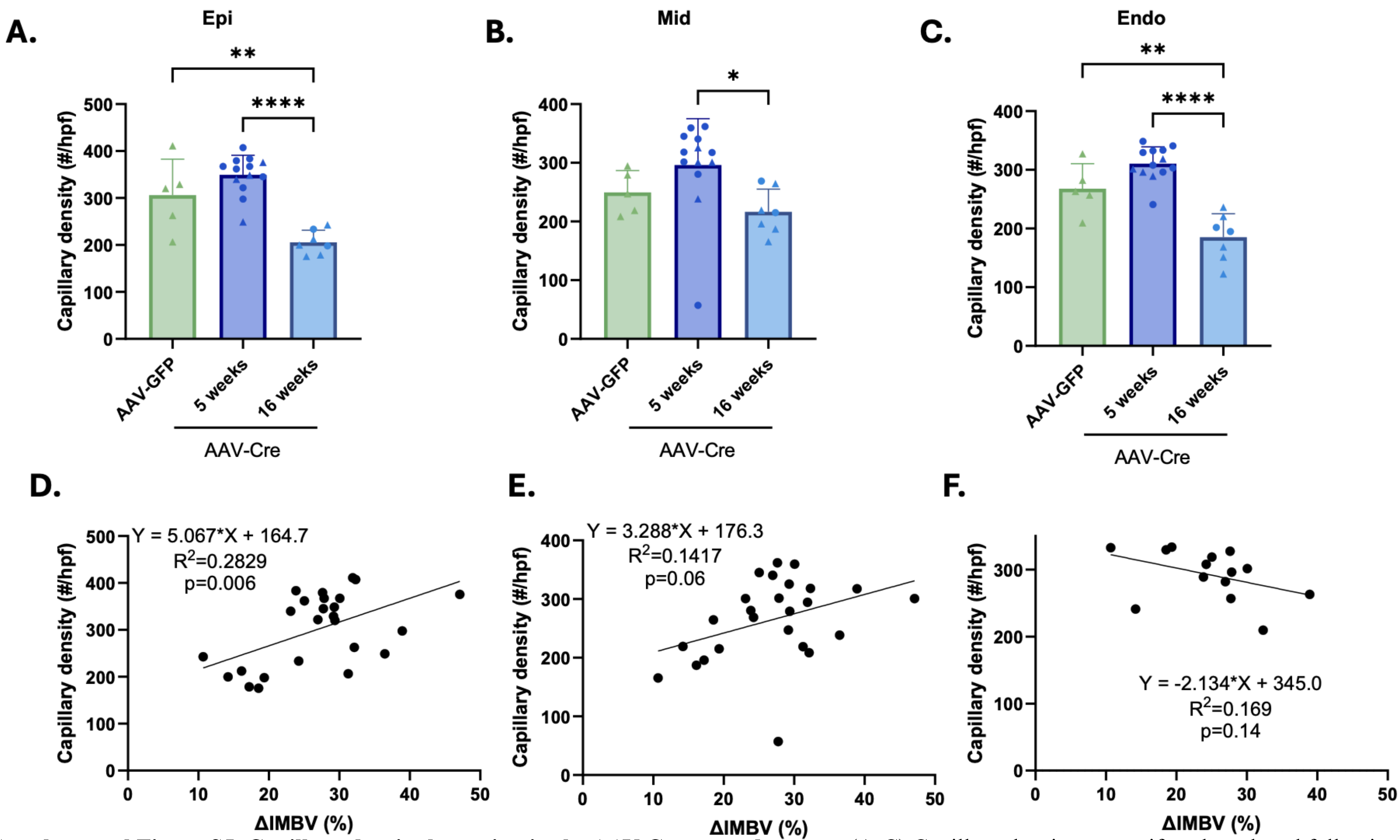


**Supplemental Figure S5. Capillary density by region in the AAV-Cre treated groups.** (A-C) Capillary density was uniformly reduced following 16 weeks of AAV-Cre expression in the epicardial (epi), mid-myocardial (mid), and endocardial (endo) regions, respectively. (D-F) Scatterplots showing the relationship between regional capillary density n the epicardial (epi), mid-myocardial (mid), and endocardial (endo) regions, respectively, and IMBV. Changes in epicardial capillary density were most strongly associated with IMBV. Data represent mean quantification of 3 images for each of 3 regions per mouse, n=5, 13, and 7per group, respectively. All groups compared by ANOVA with multiple comparisons, *$p<0.05$, **$p<0.01$, ***$p<0.001$, ****$p<0.0001$.
