## Supplemental Methods for "Linking coronary microvascular structure and function in preclinical models of coronary microvascular disease"

**Supplemental Methods for Multifractal analysis**

We used multifractal analysis to describe the space filling and spatial distribution of the coronary microvascular network. The coronary microvasculature was segmented from CD31 immunofluorescence images, converted to grayscale, binarized, and used to compute the multifractal spectrum. More specifically, key steps for multifractal analysis are as follows^1^: (1) segment the coronary microvascular from immunofluorescence images using Otsu’s method,^2^ (2) select a box shape that effectively covers the coronary microvascular structure, (3) estimate the probabilities $p_{i}\left( l \right)$ for boxes of a specific size $l$ to cover a point in the coronary microvascular structure, (4) repeat 3 for multiple box sizes exponentially distributed, (5) calculate the generalized dimension $D_{q}$, a scaled adjusted entropy defined as follows:

$$D_{q}= \frac{1}{q-1}\lim_{l\to0}\frac{ln[\sum_{i=1}^{N} p_{i}\left( l \right)^{q}]}{\ln\left( l \right)},$$

where $N$ is the number of points and $q$ acts as an adjustable magnifying glass that selects only regions with specific properties. Negative values of $q$ gives more weight to low probabilities whereas positive values give more weight to high probabilities. This allows $D_{q}$ to be adjustable, which allows for the capturing of the scaling behavior of the distribution of the coronary microvascular structure. We then (6) compute the multifractal spectrum $f\left( \alpha_{q} \right)$ by applying the Legendre transformation to $D_{q}$ to obtain $f\left( \alpha_{q} \right) = q\alpha_{q} - \left( q-1 \right)D_{q},$ where

${\alpha:=\alpha}_{q}= \frac{\partial\left[ \left( q-1 \right)D_{q} \right]}{\partial q}$is called the singularity (or Hölder)^3, 4^ and is used to capture the local scaling behavior of the coronary microvascular structure.

We use three features of the multifractal spectrum to describe the coronary microvascular: (1) the maximum value of $f(\alpha)$, termed max $f(\alpha)$; (2) argmax $f\left( \alpha\right)$, defined as the $\alpha$ value at which $f\left( \alpha\right)$ reaches its peak; and (3) the width of the multifractal spectrum. Max $f(\alpha)$is the largest fractal dimension among all the spatially differing fractal subsets of the coronary microvascular structure. Argmax $f\left( \alpha\right)$ represents the dominant local scaling behavior of the coronary microvasculature. A higher argmax $f\left( \alpha\right)$suggests a more even, spread-out network. A lower argmax $f\left( \alpha\right)$suggests that the vessels are more clumped or concentrated in some areas (i.e., not evenly distributed). High values of $\alpha$ relate to the finer branching parts of the structure. The larger the width of the multifractal spectrum, the more heterogeneous the coronary microvascular structure. We used values of max $f(\alpha)$, argmax $f\left( \alpha\right)$, and the width of the multifractal spectrum for our analyses.

3.	Balaban V, Lim S, Gupta G, Boedicker J, Bogdan P. Quantifying emergence and self-organisation of

microbial communities. Sci Rep-Uk. 2018;8. doi: ARTN 12416

10.1038/s41598-018-30654-9. PubMed PMID: WOS:000441876700119.

4.	Woods M, Bogdan P, George UZ. Polar partitioning method for the multifractal characterization of complex heterogeneous structures. Chaos Soliton Fract. 2025;201. doi: ARTN 117413

10.1016/j.chaos.2025.117413. PubMed PMID: WOS:001602528900001.
